# Female mice are more prone to develop an addictive-like phenotype for sugar consumption

**DOI:** 10.1101/2020.11.30.403535

**Authors:** Shoupeng Wei, Sarah Hertle, Rainer Spanagel, Ainhoa Bilbao

## Abstract

**Background:** The concept of “sugar addiction” is gaining increasing attention in both the lay media and scientific literature. However, the concept of sugar addiction is controversial and only a few studies have attempted to determine the “addictive” properties of sugar using rigorous scientific criteria.

**Objective:** Here we set out to systematically test the addictive properties of sugar in male and female mice using established paradigms and models from the drug addiction field.

**Methods:** Male and female C57BL/6N (8-10 weeks old) were evaluated in 4 experimental procedures to study the addictive properties of sugar: (i) a drinking in the dark (DID) procedure to model sugar binging; (ii) a long-term free choice home cage drinking procedure measuring the sugar deprivation effect (SDE) following an abstinence phase; (iii) a long-term operant sugar self-administration with persistence, motivation and compulsivity measures and (iv) intracranial self-administration (ICSS).

**Results:** Female mice were more vulnerable to the addictive properties of sugar than male mice, showing higher binge and long-term, excessive drinking, a more pronounced relapse-like drinking following deprivation, and higher persistence and motivation for sugar. No sex differences were seen in a compulsivity test or reward sensitivity measured using ICSS following extended sugar consumption.

**Conclusion:** This study demonstrates the occurrence of an addictive-like phenotype for sugar in male and female mice, similar to drugs of abuse, and suggests sex-dependent differences in the development of sugar addiction.

## Introduction

The terms substance use and addictive behaviors encompass disorders that develop as a result of the use of psychoactive substances. However, the term addiction is increasingly also applied to a range of other problematic behaviors, such as excessive gaming behavior. Consequently, the new psychiatric classification systems ICD-11 has now categorized drug addictions *vs*. so-called behavioral addictions (Reed et al., 2019). This marks a milestone in diagnosis and psychiatry. Despite the introduction of newly introduced behavioral addictions into ICD-11, there is public awareness that other behaviors such as excessive food/sugar consumption (Wiss et al., 2018) can lead to addiction as well. However, this remains debatable from a scientific perspective; it has been proposed that non ICD-11 classified behavioral addictions should be classified using terms such as “problematic use” and described in their sociocultural context rather than within an addiction framework (Wiss et al., 2018).

Although there is no consensus on the behavioral construct of “sugar addiction,” it is assumed that its problematic use, similar to chronic drug abuse, can lead to a complex behavioral disorder involving multiple interactions between genetic, biological and environmental aspects. In fact, the behavioral phenotypes associated with chronic drug abuse and problematic sugar consumption are similar with respect to compulsive overconsumption, craving, loss of control, and even withdrawal responses (Wiss et al., 2018; Avena et al., 2009). It has been proposed that specific foods, especially those that are rich in sugar, are capable of promoting addiction-like behavior and neuronal change only under certain conditions. That is, these highly sugar-containing foods, although highly palatable, are not addictive *per se* but become so following a restriction/binge pattern of consumption (Corwin and Grigson, 2009; Pelchat, 2009). Importantly, alcohol or drugs of abuse are also not addictive *per se* but become so following a restriction/binge pattern of consumption (Foo et al., 2017; Spanagel, 2017). Thus, a critical factor in the development of alcohol or drug addiction is restriction/deprivation and multiple phases of deprivation and binging that can lead, at least in some individuals, to addictive behavior (Foo et al., 2017; Spanagel, 2017).

Epidemiological studies have observed significant gender-specific differences in patients with addiction disorders, with females more vulnerable to the initiation of drug use and the development of excessive chronic and even compulsive behavior (Hudson and Stamp, 2011; Becker et al., 2016). Similarly, women are more inclined to the excessive intake of sugar and a greater incidence of obesity (Bennet et al., 2018; Ogden et al., 2015). Similar to humans, extensive preclinical research with laboratory animals has found sex differences across different drug-related addictive phenotypes and mechanisms, revealing that females are more vulnerable to the initiation of drug use and subsequent withdrawal and relapse stages (Becker et al., 2016; Sanchis-Segura and Becker, 2016). While fewer studies have evaluated sex differences in food/sugar-related phenotypes, one hypothesis is that sex differences related to food addiction may occur in rodents and parallel those in humans (Becker et al., 2016; Sample and Davidson, 2018).

There have been several attempts to model different aspects of excessive sugar consumption in rodents. For example, repeated excessive intake of sugar created a state in which the administration of an opioid antagonist resulted in behavioral and neurochemical signs of withdrawal (Colantuoni et al., 2002). Furthermore, excessive sugar intake cross-sensitized to amphetamine in rats (Avena and Hoebel, 2003). The repeated, excessive intake of sugar can also lead - following deprivation - to a sugar deprivation effect (SDE) (Avena et al., 2005), which is considered a relapse-like phenomenon in the addiction field (Spanagel, 2017; Vengeliene et al., 2014). Several more aspects of repeated, excessive intake of sugar on behavioral and cognitive function (Kendig, 2014) have been studied in rodents; however, a comparative study on sex differences in different paradigms and animal models used in the addiction field has so far not been conducted.

Here we systematically tested the addictive properties of sugar in male and female mice using established paradigms and models from the drug addiction field. (i) We applied the drinking in the dark (DID) procedure to model sugar binging. The DID procedure was originally developed to induce alcohol intoxication and model alcohol binging in mice (Rhodes et al., 2005; Thiele and Navarro, 2014). (ii) We also used a long-term free choice home cage drinking procedure and studied the sugar deprivation effect (SDE) following an abstinence phase. The deprivation effect is a measure of consumption during a relapse-like situation in the addiction field (Spanagel, 2017; Vengeliene et al., 2014). (iii) We also introduced a long-term operant sugar self-administration paradigm and quantified three criteria for addictive behavior: persistence, motivation and compulsivity. These three criteria have been used to operationalize cocaine addiction in rats (Deroche-Gamonet and Piazza, 2014) and are now increasingly applied to other addictions in preclinical research (Spanagel, 2017; Mancino et al., 2015).

## Methods

### Animals

For all four experiments, we used 8-10 weeks-old C57BL/6N mice. Mice were single-housed in standard hanging cages at 21 ±1°C and 50±5% relative humidity on a reversed 12h light/dark cycle, with lights on at 7:30p.m. The animals were provided with standard rodent food (Altromin Spezialfutter GmbH & Co, LASQCdiet Rod16-H. Composition: cereals, vegetable by-products, minerals, oils and fats, yeast; crude nutrients: 16.30 % crude protein, 4.30% crude fat, 4.30% crude fibre, 7.00% crude ash), a bottle containing 5% (w/v) sugar solution during the DID and long-term sugar paradigms (see below for details) and tap water *ad libitum*. All the experiments were performed in the dark cycle. All mice were handled on a daily basis before starting the experiments and were habituated to the behavioral testing environments. Procedures for this study complied with the regulations covering animal experimentation within the European Union (European Communities Council Directive 86/609/EEC) and Germany (Deutsches Tierschutzgesetz) and the experiment was approved by the German animal welfare authorities (Regierungspräsidium Karlsruhe).

### Experiment 1. Binge sugar drinking: Drinking in the Dark (DID) paradigm

For the DID paradigm, 22 male and 19 female mice were used. We used a modified version of the standard DID protocol originally developed by Rhodes et al. (2005) to model binge consumption of sugar. In brief, during the first 3 days, 4 h into the dark cycle, animals had access for 2 hours to standard food and a single bottle containing 5% (w/v) sugar solution and on the fourth day, mice were exposed for 4 hours under identical conditions (food and sugar). Sugar solution was freshly prepared daily, and sugar consumption was calculated using the Drinkometer system (see below). Data collected during the 4-h DID period were expressed as g/kg body weight for sugar intake. After the DID, all animals had 24h access to sugar during the next 3 days and the amount consumed during both conditions was used for statistical analysis.

### Experiment 2. Home cage two-bottle free choice sugar drinking and assessment of relapse-like drinking by means of the sugar deprivation effect (SDE)

For this experiment, 42 male and 41 female mice were used. Mice had continuous free choice access to a bottle containing a sugar solution (5% w/v) solution and a bottle with tap water in the homecage for 8 weeks. During the first and last 3 days of sugar exposure, sugar and water intake was recorded using the Drinkometer system (Eisenhardt et al., 2015a, 2015b; Bilbao et al., 2019). Mice were afterwards deprived from sugar for 12–15 days, during which they only had access to two bottles of tap water. After the deprivation period, the SDE was tested for 72 h by reintroducing the sugar bottle.

Sugar (g/kg) and water (ml) intake as well as the sugar preference (% of total fluid intake) were calculated per day. During baseline and SDE measurements, sugar and water intake was additionally calculated in 4 h time intervals. Baseline sugar and water intake was calculated as the mean of the last 3 d of baseline recording.

### Experiment 3. Long-term operant self-administration and assessment of motivation, persistence and compulsivity for sugar consumption

Twenty-three male and 23 female C57BL/6N mice were used as for the operant sugar self-administration. Mice were trained and tested in operant chambers (TSE Systems, Bad Homburg, Germany). Each chamber had two ultrasensitive levers (required force, ≤1 g) on opposite sides: one functioning as the active and one as the inactive lever. Next to each lever, a front panel containing the visual stimulus was installed above a drinking microreservoir. When the programmed ratio requirements were met on the active lever, 10 μl of the 10% sugar solution was delivered into a microreservoir, and the visual stimulus was presented via a light located on the front panel. Responses on the inactive lever were recorded but had no programmed consequences.

Mice were trained to self-administer 10% sugar (w/v) in 30 min daily sessions on a fixed ratio (FR) 1 schedule of reinforcement, where each response resulted in delivery of 10 μl of fluid. Following a lever press a 5 s time-out period was in effect, during which responses were recorded but not reinforced. After 15 sessions the response requirements were enhanced to a FR4 schedule. After 30 sessions under FR4 responding and the establishment of stable baseline responding mice were tested for an addictive-like phenotype.

At the end of the training we measured first the persistence, then the motivation, and finally the compulsivity for sugar self-administration. The persistence of response was measured using the time out (TO) test as described previously (Cannella et al., 2013, 2018). This 30-min session was composed of two 12-min available periods separated by one 6-min unavailable period as “time-out’during which the house light went on and active pressings resulted in no consequences except being registered. The tests were repeated for another three times, of which the first session was considered as the habituation to novelty, and the mean value of the last three sessions were calculated as a measure for persistence in responding for sugar when sugar was not available. Motivation for sugar was measured by a progressive ratio (PR) schedule of reinforcement in which the response requirement (the number of lever responses required to receive the reward) was progressively increased: the reinforcement schedule increased consecutively from ratio 1 by step size 2 for each reward. The last accomplished ratio was taken as the breaking point. Sessions were ended when mice were not earning a reward for 30 min. Compulsive-like behavior or resistance to aversive stimuli was measured by taste adulteration with quinine. A 0.8mM quinine concentration was added to the sugar solution and the percentage of reduction over baseline was taken as a measure of compulsivity. Between the different test situations, mice were measured for 3-4 days for their baseline self-administration.

### Experiment 4. Assessment of brain stimulation reward by intracranial self-stimulation (ICSS)

The last experiment involved 6 male and 8 female sugar experienced mice. In order to assess the function of the brain reward system, ICSS was performed as described previously (Bilbao et al., 2015, 2020) in a subset of mice from experiment 2 (continuous free choice access to a bottle containing a sugar solution (5% w/v) solution and a bottle with tap water in the home cage for 8 weeks). Briefly, mice were anesthetized with 1.5-1.8% of isoflurane (CP-Pharma, Burgdorf, Germany), stereotaxically implanted with insulated monopolar stainless steel electrodes (.28 mm diameter) (Plastics One, USA) to the right medial forebrain bundle in the lateral hypothalamus (coordinates from Bregma: anterior (AP) −1.2, lateral (ML) +1, ventral (DV)–5.4), and trained to respond for brain stimulation reward (BSR).

During each testing session, mice responded during three consecutive series of 15 descending frequencies (.05 log10 steps). Maximum control rate (MCR), and total number of stimulations were calculated from the mean of the second and third series. Stimulation seeking and extinction components were calculated from the total number of stimulations during the first 5 highest frequencies and the remaining 10, respectively.

### Statistics

Statistical analyses were performed by ANOVA with Newman–Keuls test for *post hoc* comparisons using Statistica 10 (StatSoft). All values are given as mean ± SEM, and statistical significance was set at *p* < 0.05.

## Results

### Experiment 1. Characterization of binge sugar drinking in male and female mice using the DID paradigm

Here we defined sugar binge drinking as the intake of an excessive amount of sugar in a short period. Thus, the amount of restricted sugar intake (in g/kg) consumed during the first 4 hours on the fourth day into the dark period exceeded significantly sugar consumption when measured 4 hours into the dark period in the same animals when they had unrestricted free choice sugar consumption (Fig. 1A, Two-way ANOVA, *Exposure* effect: F_(1,39)_ =31.4; *P*<0.0001; *Sex* effect: F_(1,39)_ =2.7; *P*=0.1). The increase in females (74% over continuous) was greater than in males (56% over continuous). Because there was a main effect of exposure, but not a significant exposure x sex interaction, we performed a one-way ANOVA by sex. Indeed, although continuous free choice access did not differ between both sexes (F_(1,39)_ =1; *P*=0.3), females showed a significantly higher intake during binge exposure compared to males (F_(1,39)_ =4.3; *P*<0.05).

**Figure 1.**
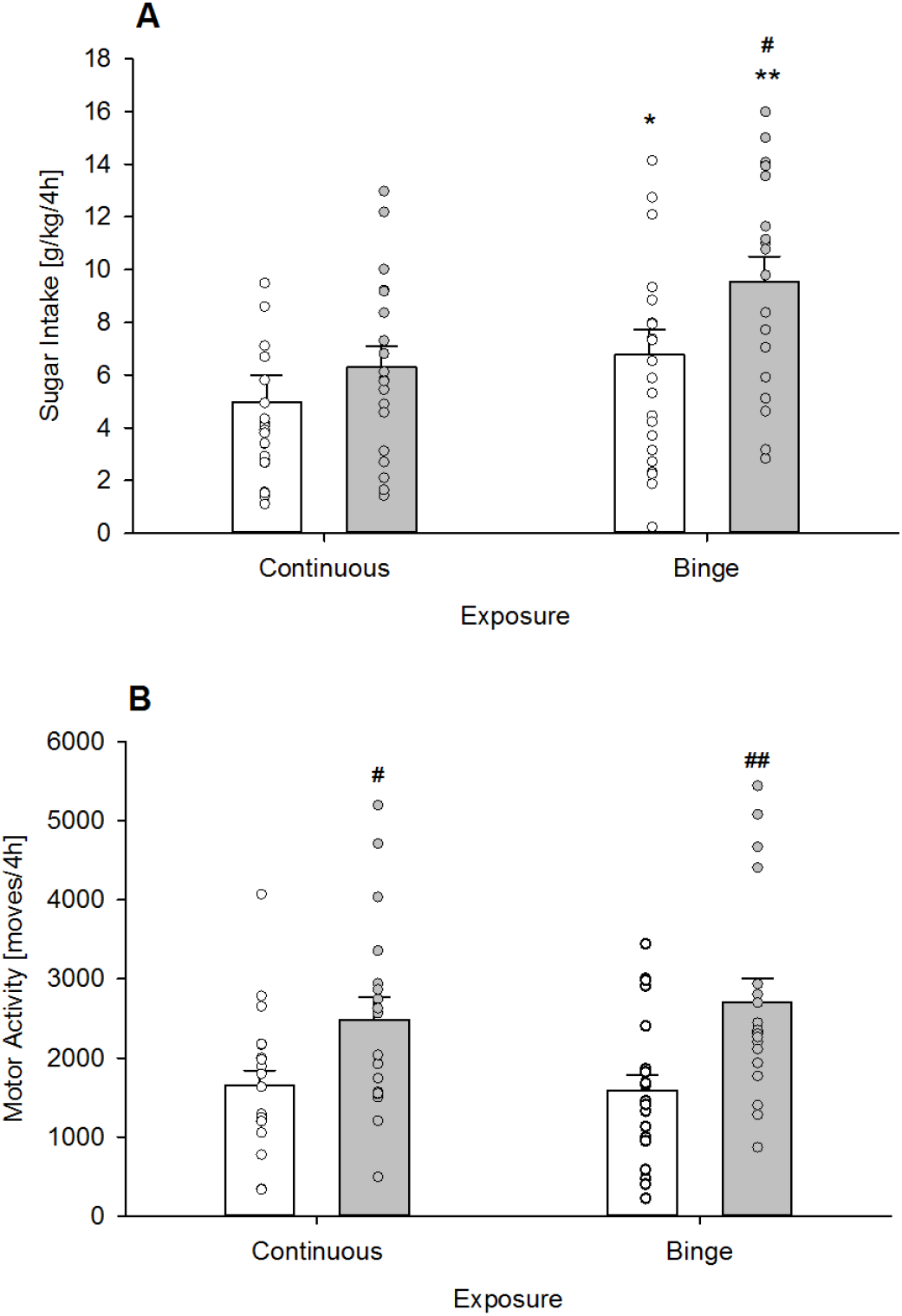
Characterization of sugar binge drinking and locomotor activity in male and female mice (Experiment 1). (A) During binge exposure (restricted sugar consumption during the first 4h into the night active period) sugar intake was significantly higher in male (n=22) and female (n=19) mice compared to a continuous free choice access. (B) Locomotor activity did not differ between binge and continuous sugar exposure. However, females displayed a general increased locomotion compared to males. All data represent mean + SEM. (*), (**) indicate *P*< 0.01 and 0.001 versus continuous exposure, respectively; (#), (##) indicate *P*< 0.05 and 0.01 versus male mice, respectively.

When analyzing the locomotor activity, we found that binge sugar access did not have any effect (Fig. 1B, *Exposure* effect: F_(1,39)_ =0.2; *P*=0.7). However, females displayed a general increased locomotion (34% over continuous) when compared to male (8% over continuous) mice (Fig. 1B, *Sex* effect: F_(1,39)_ =11.9; *P*<0.05).

### Experiment 2a. Characterization of acquisition and maintenance of sugar drinking in male and female mice

During the acquisition phase (the first 3 days of free-choice sugar drinking), both males and females showed diurnal rhythmicity, with higher drinking levels during the dark, active phase compared with the light, inactive phase of the day (Fig. 2A, Fig. 2B, two-way ANOVA, *Phase* effect: F_(5 405)_ =77; *P*<0.001) with no differences between males and females (*Sex* effect: F_(1,81)_ =0.3; *P*=0.5). Consequently, the total sugar intake during 24h was similar in both groups (Fig. 2C, One-way ANOVA, F_(1,81)_ =2.6; *P*=0.1). Diurnal rhythmicity was not that marked in the daily preference over water (Fig. 2D). However, the mean of the first 3 days indicated a diurnal pattern (Fig. 2E, *Phase* effect: F_(5,405)_ =10.4; *P*<0.001), with females showing higher preference than males between the end of the active and beginning of the inactive phase (*Sex* effect: F_(1,81)_ =10.6; *P*<0.01; *Sex x Phase* interaction effect: F_(5,405)_ =2.8; *P*=0.05), which resulted in a higher total preference (94%) compared to males (90%) (Fig. 2F, *t*_(81)_=-3.2; *P*<0.01). Similar to the intake pattern, locomotor activity (Fig. 2G) also showed typical diurnal rhythmicity (Fig. 2H, *Phase* effect: F_(5,405)_ =40.5; *P*<0.001) and females showed higher locomotor activity (*Sex* effect: F_(1,81)_ =9.6; *P*<0.01) which was restricted to the active phase (*Phase x Sex* interaction effect: F_(5,405)_ =5.2; *P*=0.001). As a result, the total daily locomotor activity was increased in females compared to males (Fig. 2I, F_(1,81)_ =5.9; *P*<0.05).

**Figure 2.**
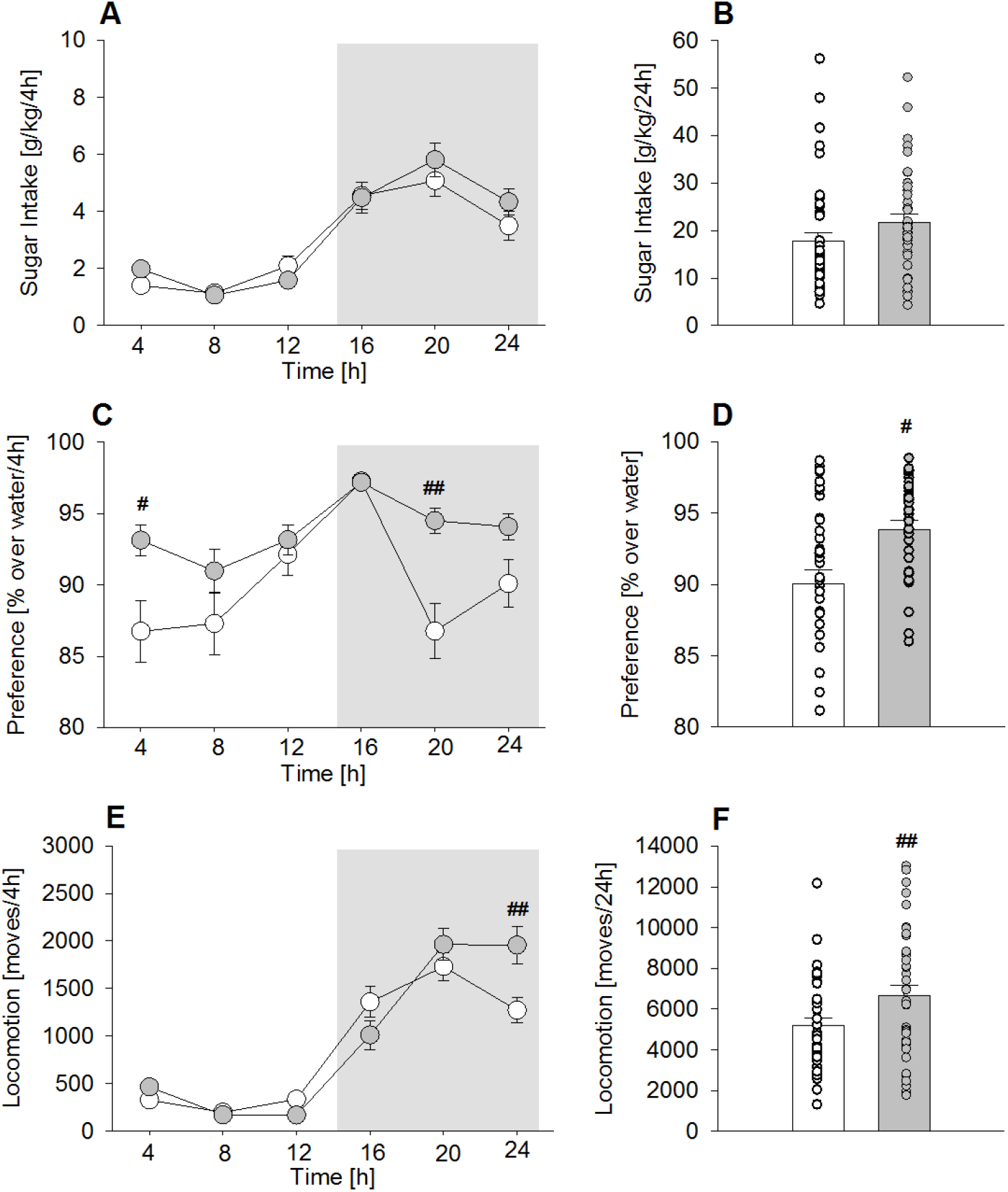
Acquisition of sugar consumption in male (n=42) and female (n=41) mice (Experiment 2). Sugar drinking (A-B), preference over water (C-D) and locomotor activity (E-F) during the first 3 days of free choice sugar exposure. (A) Both males and females showed the typical diurnal rhythmicity, with higher drinking levels during the dark, active phase compared with the light, inactive phase of the day with no differences between males and females (all day points differed significantly from all night points, not indicated). (B) Consequently, the total sugar intake during 24h was similar in both groups. (C) Such rhythmicity was not that marked in the daily preference over water but the mean showed a clear diurnal pattern, with females showing higher preference between the end of the active and beginning of the inactive phase, (D) which resulted in a higher total preference compared to males. (E) Similar to the intake pattern, locomotor activity also showed the typical diurnal rhythmicity (all day points differed significantly from all night points, not indicated). Females showed higher locomotor activity which was restricted to the active phase. (F) As a result, the total daily locomotor activity was higher in females than in males All data represent means ± SEM. (#), (##) indicate *P*< 0.05 and 0.001 versus male mice, respectively.

After a long-term drinking period of 8 weeks, the diurnal rhythmicity of sugar intake remained typical (Fig. 3A; Fig. 3B, *Phase* effect: F_(5,405)_ =71.7; *P*<0.001), but female mice showed higher intake during the active phase (Fig. 3B, *Phase x Sex* interaction: F_(5,405)_ =6.1; *P*<0.001). In the total 24h intake, females consumed significantly more (53%) than males (Fig. 3C, F_(1,81)_ =8.1; *P*<0.01). In contrast to the pattern displayed during the acquisition phase, the daily preference after long-term drinking had clearly a diurnal pattern (Fig. 3D; Fig. 3E, *Phase* effect: F_(5,405)_ =36.6; *P*<0.001), which did not differ between both groups (*Sex* effect: F_(1,81)_ =1.1; *P*=0.3). Total 24h preference was also not different (Fig. 3F, F_(1,81)_ =1.1; *P*=0.3). The locomotor activity (Fig. 3G) remained high in the females and similar to the acquisition phase, restricted to the active phase (Fig. 3H, *Phase*: F_(5,405)_ =64; *P*<0.0001, *Gender*: F_(1,81)_ =0.6; *P*=0.4).This increase did reach significance when considering the total 24h activity (Fig. 3I, F_(1,81)_ =3.9; *P*=0.05).

**Figure 3.**
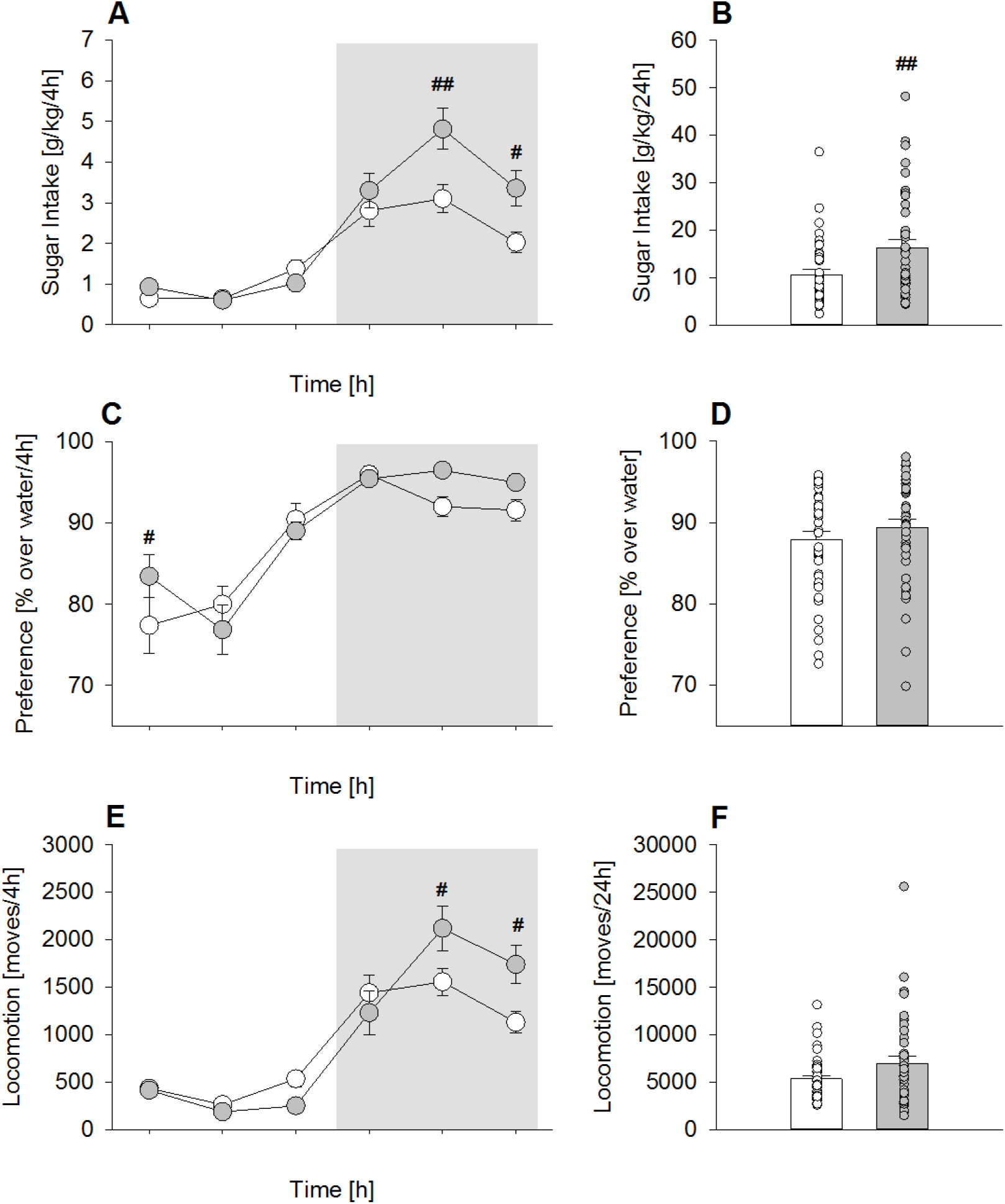
Chronic long-term sugar drinking in male (n=42) and female (n=41) mice (Experiment 2). Sugar drinking (A-B), preference over water (C-D) and locomotor activity (E-F) during the last 3 days of a 6 weeks period of free choice sugar exposure. (A) There was a clear diurnal rhythmicity of sugar intake, but the females showed higher intake during the active phase. (B) As shown for 24h intake, females consumed significantly more sugar than males. (C) The daily preference after long-term drinking showed a diurnal pattern and the mean taken of the last 3 days did not differ between both groups; total 24h preference was also not different (D). (E) The locomotor activity remained high in the females and, similar to the acquisition phase, was restricted to the active phase. (F) However, the total daily locomotor activity did not differ between females and males. All data represent means ± SEM. (#), (##) indicate *P*< 0.05 and 0.01 versus male mice, respectively.

Importantly, chronic sugar intake over 8 weeks led to a significant increase in body weight when compared to the sugar-free period, one week before (Fig. 4A, Two-way ANOVA; Fig. 4A, B *Sugar* effect: F(1,27) =31.1; *P*<0.001; *Time* effect: F(7,189) =61.5; *P*<0.001) but no sex differences were observed (*Sex* effect: F(1,27) =0.35; *P*=0.56). After 8 weeks, the total body weight increase was similar in males and females (Fig. 4B, *Sex* effect: F(1,27) =0.5; *P*=0.5), and significantly increased compared to the control group (*Sugar* effect: F(1,27) =55.7; *P*<0.001).

**Figure 4.**
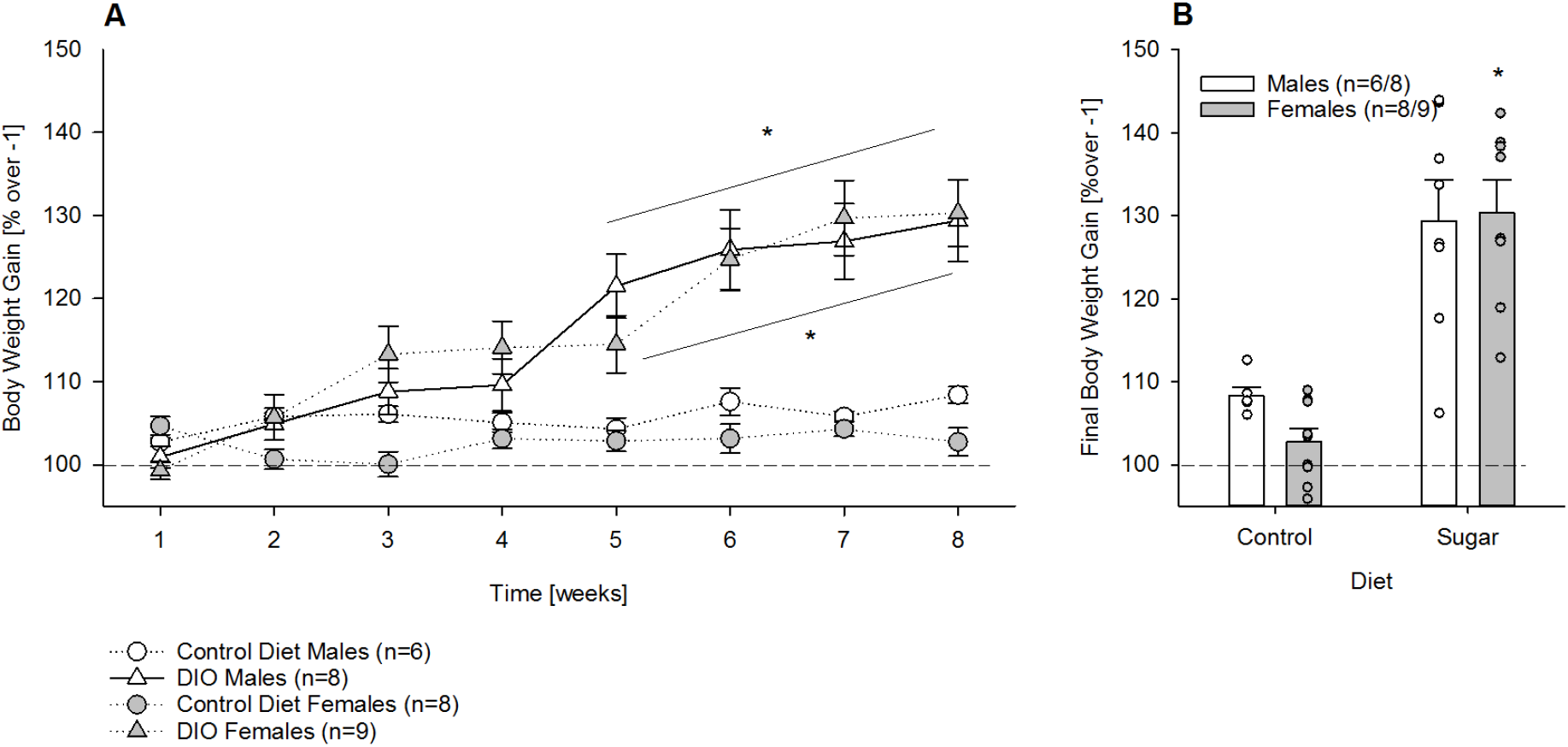
Pronounced increase in bodyweight during chronic long-term sugar drinking in male (n=6/8) and female (n=8/9) mice (Experiment 2). (A) Compared to the sugar-free week before (week −1), chronic sugar intake over 8 weeks led to a significantly higher increase in bodyweight than in the water drinking control group but no sex differences were observed. (B) After 8 weeks, the total body weight increase was similar in males and females and significantly increased compared to the control group. All data represent means ± SEM. (*) indicate *P*< 0.001 versus water control group.

### Experiment 2b. Characterization of relapse-like sugar intake (sugar deprivation effect (SDE) in male and female mice

After 8 weeks of home cage voluntary sugar consumption all animals underwent a 15 days period of sugar deprivation which significantly increased the 24h sugar intake compared to the baseline (mean of the last 3 days) in both male and female mice, indicative of a sugar deprivation effect (SDE, Fig. 5), which lasted two days (Fig. 5A, two-way ANOVA; *Deprivation* effect: F_(3,243)_ =23.9; *P*<0.0001), and was higher in females (*Sex* effect: F_(1,81)_ =6.9; *P*<0.05). During the first day, the division of the SDE into 4h intervals showed that the SDE was strongly pronounced during the first 4h of sugar re-exposure and lasted not longer than 8h in both males and females (Fig. 5B, *Deprivation* effect: F_(23,1863)_ =74; *P*<0.0001). However, during the second day the SDE was stronger in females compared to males (*Sex* effect: F_(1,81)_ =6.8; *P*<0.05), as it lasted again 8h, while in males it was attenuated in respect to the first day and lasted only 4h (*Sex x Deprivation* effect: F_(3,243)_ =7.7; *P*<0.0001).

**Figure 5.**
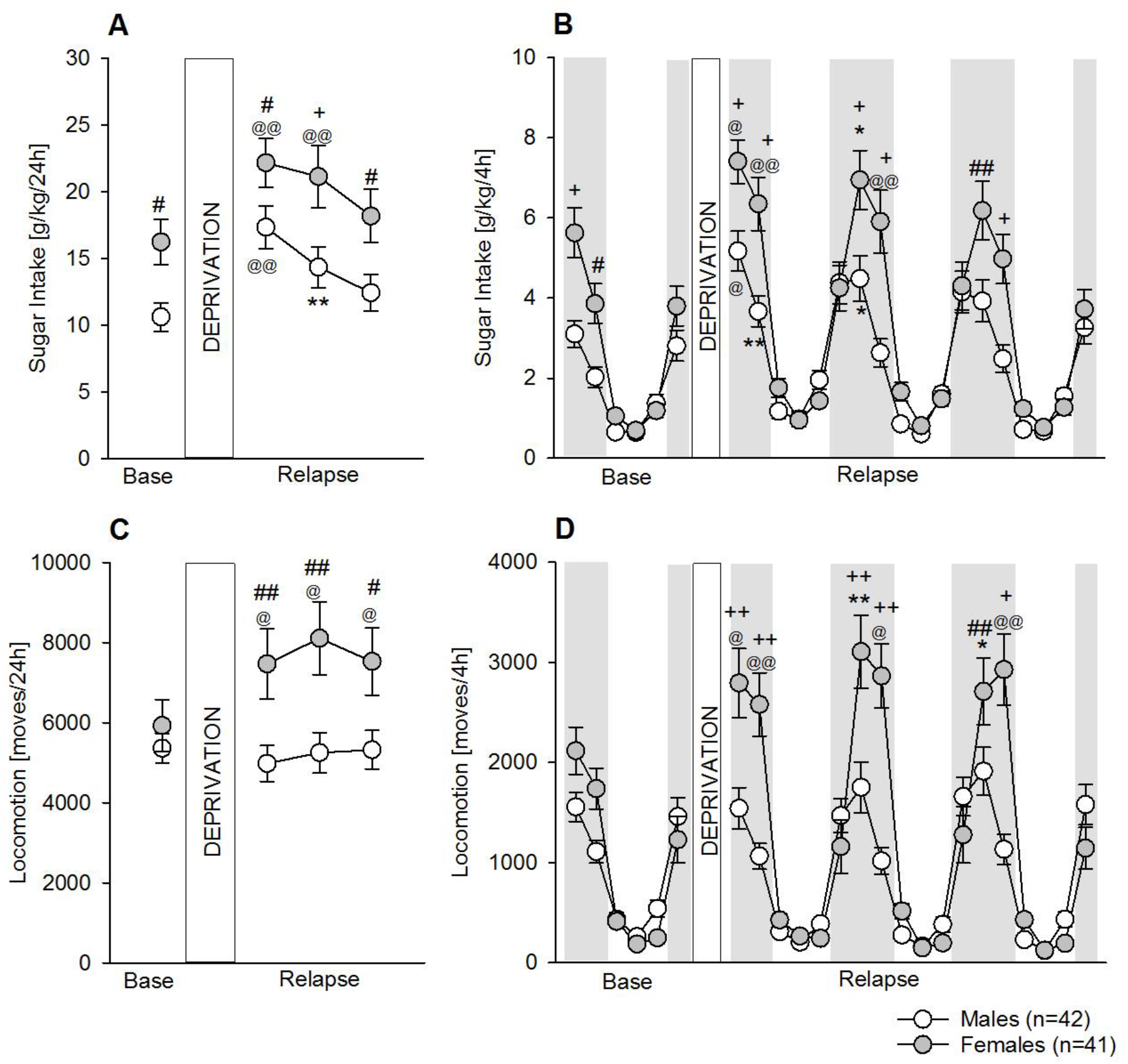
Characterization of the sugar deprivation effect (SDE) in male and female mice (Experiment 2). (A) A period of sugar deprivation significantly increased the sugar intake in both male (n=42) and female (n=41) mice, indicative of a sugar deprivation effect (SDE), which lasted 2 days, and was higher in females. (B) During the first day, dissection of the SDE into 4h intervals showed that intake was strongly pronounced during the first 4 hours of sugar re-exposure and lasted for 8 hours in both males and females. During the second day, the SDE was stronger in females compared to males, as it lasted for 8 hours, while in males it was attenuated in respect to the first day and lasted only 4 hours. (C) Females (but not males) showed a very pronounced increase in locomotor activity which lasted 3 days. (D) Dissection of the locomotor activity into 4h intervals showed that it was strongly pronounced during the first 4 hours of sugar re-exposure and lasted not longer than 8 hours in females. All data represent means ± SEM. +,++) indicate *P*< 0.05; 0.01; 0.005; 0.001 vs male mice; respectively; (*, *, @,@@) indicate *P*< 0.05; 0.01; 0.005; 0.001 vs corresponding baseline point, respectively.

Locomotor activity analysis also showed a very pronounced increase in females which lasted 3 days (Fig. 5C). This increase was not only significant compared to baseline (*Deprivation* effect: F_(3,243)_ =4.1; *P*<0.01), but also strongly differed across sex (*Sex* effect: F_(1,81)_ =5.9; *P*<0.05; *Sex x Deprivation* interaction effect: F_(3,243)_ =5.4; *P*<0.01). Almost identical to the intake pattern, division of the locomotor activity into 4h intervals showed that it was strongly pronounced during the first 4h of sugar re-exposure (Fig. 5D, *Deprivation* effect: F_(23,1863)_ =63.3; *P*<0.0001) and lasted not longer than 8h in females (Fig. 5D, *Sex*: F_(1,81)_ =5.8; *P*<0.05 and *Sex x Deprivation*: F_(3,243)_ =12.2; *P*<0.0001 effects).

### Experiment 3a. Characterization of long-term operant self-administration of sugar in male and female mice

A third group of mice was trained to self-administer sugar. During the FR1 phase, all mice similarly acquired and maintained stable active lever pressing for sugar (Fig. 6A), and the mean from the last 5 sessions did not differ between male and female mice (Fig. 6B, F_(1,44)_ =1.6; *P*=0.2). However, as observed in Fig. 6C, inactive lever pressing was higher in female mice across the 15 FR1 sessions, and indeed the mean of the last 5 sessions showed significant differences compared to male mice (Fig. 6D; F_(1,44)_ =6.2; *P*<0.05). During the FR4 phase, the overall performance of the females was higher than males: females earned more rewards and made more responses on the inactive lever (Fig. 6A,C); however, averaging the last 5 days did not result in significant differences (only a trend) between sexes (Fig. 6B: F_(1,44)_ =2.8; *P*=0.1; and Fig. 6D: F_(1,44)_ =2.4; *P*=0.1).

**Figure 6.**
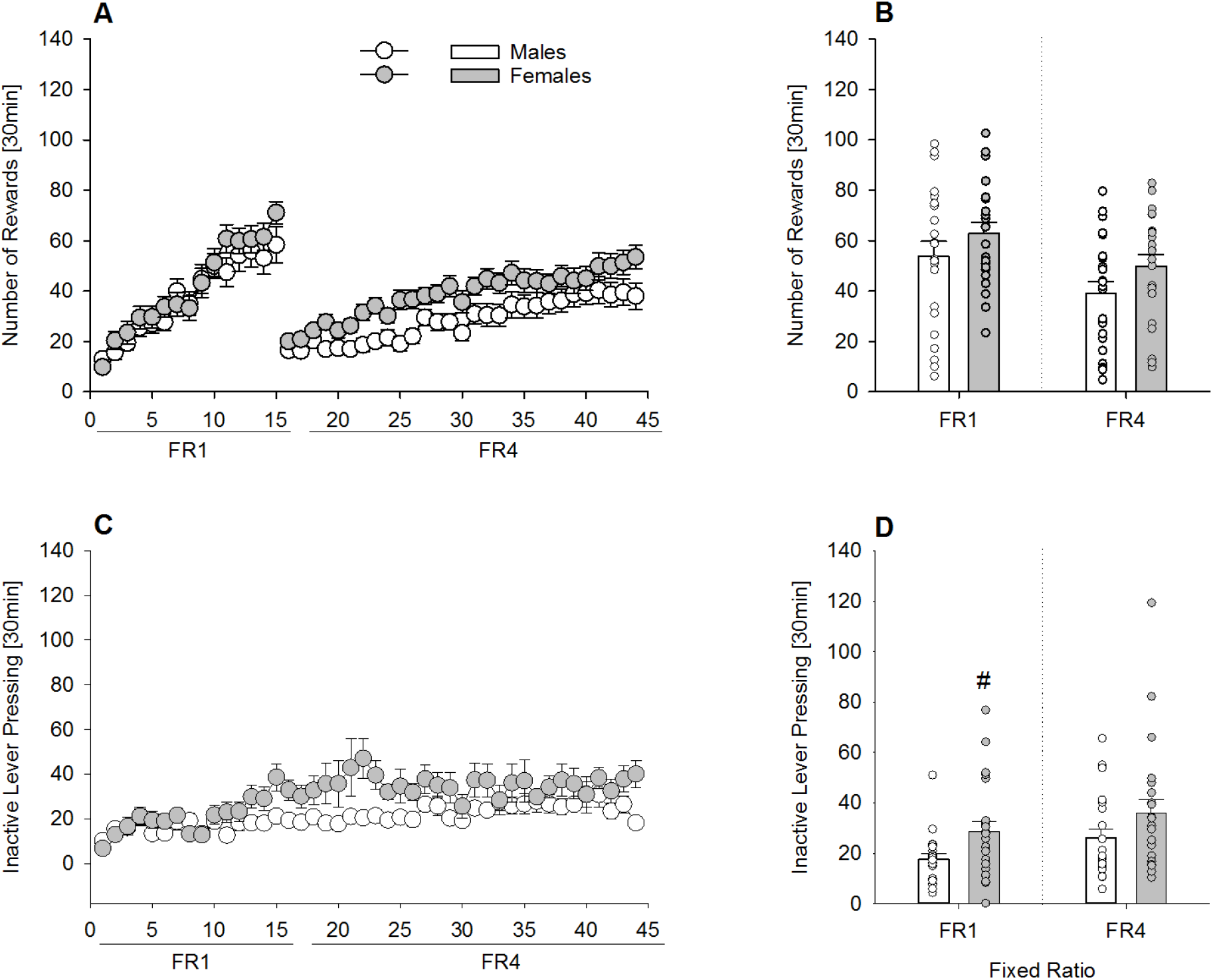
Long-term operant self-administration for sugar in male (n=23) and female (n=23) mice (Experiment 3). (A) Number of rewards earned, (C) inactive lever pressing per sessions, during FR1 (15 sessions) and FR4 (29 sessions) schedule of reinforcement. Mean from the last 5 sessions of (B) rewards earned, and (D) inactive lever pressing. (A) During the FR1 phase, all mice similarly acquired and maintained stable active lever pressing for sugar and (B) the mean from the last 5 sessions did not differ between male and female mice. (C) Inactive lever pressing was higher in female mice across the 15 FR1 sessions, and (D) the mean of the last 5 sessions showed a significant increase compared to male mice. During the FR4 phase, (A) females earned more rewards and made more responses on the inactive lever (C, D). All data represent mean ± SEM. # indicates *P*<0.05 versus male mice.

### Experiment 3b. Characterization of an addiction-like phenotype for sugar in male and female mice

Next, the addictive-like phenotypes of persistence, motivation, and compulsivity were studied using the time out (TO), progressive ratio (PR), and quinine test, respectively (Fig. 7). During the TO test, the number of lever presses per min was almost doubled in females compared to male mice (Fig. 7A, F_(1,44)_ =8.8; *P*<0.005) indicating higher persistence for sugar-seeking. Motivation was assessed by a PR schedule of reinforcement. As illustrated in Fig. 7B, the breakpoint that male mice were willing to obtain sugar was around 17, while in females it was significantly increased to almost 23 (F_(1,44)_ =5; *P*<0.05), indicating a higher motivation. The last test measured compulsive-like behavior by taste adulteration with quinine (Fig. 7C). A 0.8mM quinine concentration added to the sugar solution reduced the lever pressing to obtain sugar similarly in both male (79%) and female (71%) mice, and no significant differences were found (F_(1,44)_ =1.5; *P*=0.2). Thus, in contrast to persistence and motivation, compulsive-like behavior for sugar was not higher in females compared to male mice.

**Figure 7.**
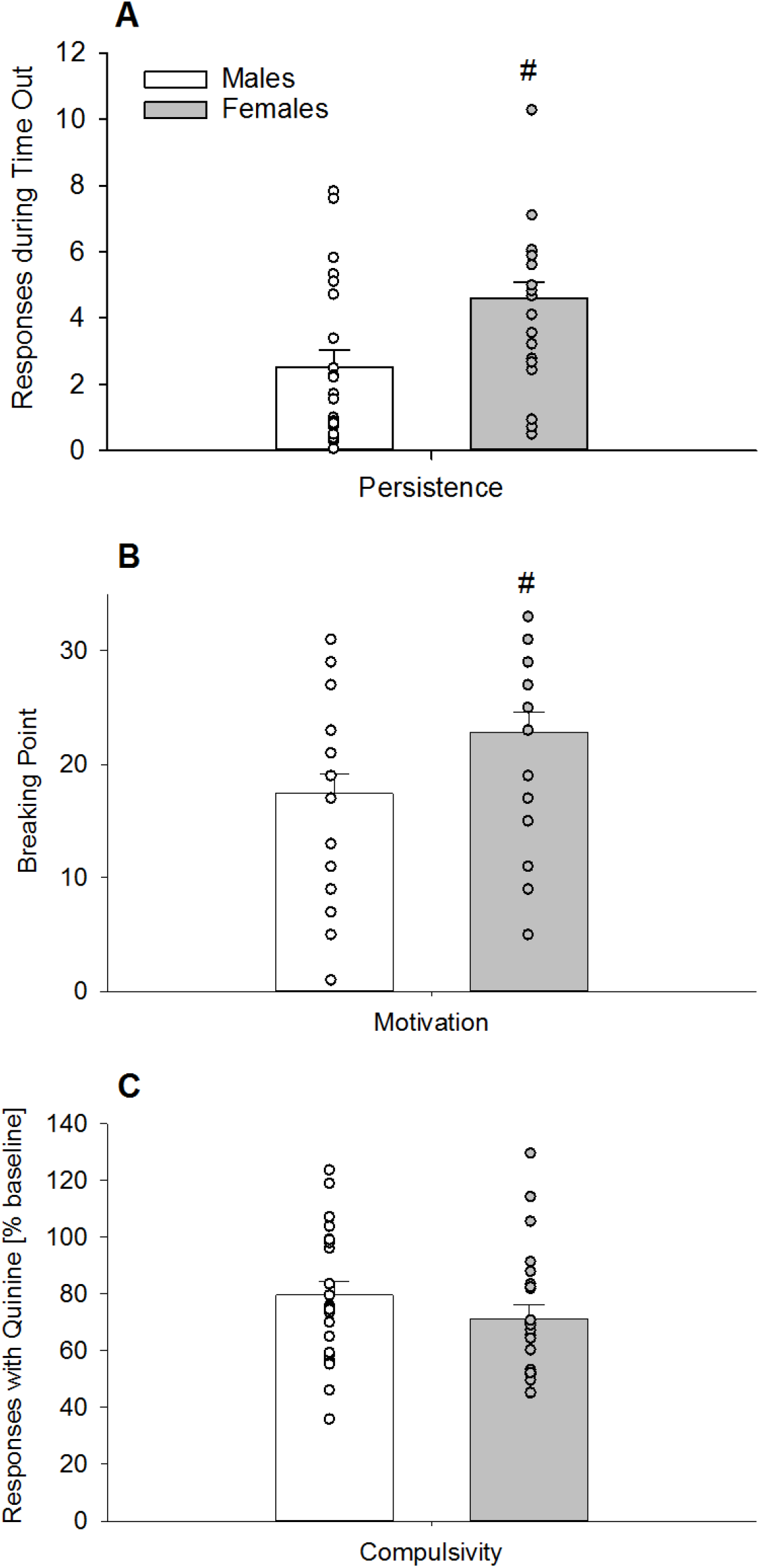
Addictive-like phenotypes of persistence, motivation and compulsivity in long-term operant sugar self-administering male (n=23) and female (n=23) mice (Experiment 3). (A) Persistence was measure by the TO test. During the single 6 min time out of sugar access, the number of lever presses per min was almost doubled in females compared to male. (B) Motivation as assessed by a PR schedule of reinforcement. The breakpoint or limit to the amount lever pressing that male mice were willing to perform to obtain sugar was around 17, while in females it was significantly increased to almost 23. (C) Compulsive-like behavior by taste adulteration with quinine reduced the number of lever presses to obtain sugar similarly in both male and female mice. All data represent mean + SEM. (#) and (##) indicate *P*<0.05 and *P*< 0.01 versus male mice, respectively.

### Experiment 4. Characterization of reward sensitivity by ICSS in male and female mice

We further tested a fourth group of male and female mice brain stimulation reward using the ICSS paradigm (Bilbao et al., 2015, 2020). All mice showed the expected frequency-dependent decrease in the number of stimulations per trial which became significant from the 6^th^ frequency on (Fig. 8A, two-way ANOVA, *frequency* effect: F_(14,168)_ =26.8; *P* <0.0001). During the “seeking” and “extinction” components the performance of both male and female mice was almost identical (Fig. 8B, *Sex* effect (_F(1,12)_ =0.00; *P* =1), and was characterized by a significant 40% reduction during the “extinction” component compared to the “seeking” component (*Component* effect (F_(1,12)_ =16; *P* <0.01). In line with this, all mice also showed an indistinguishable reinforcement rate (Fig. 8C, F_(1,12)_ =0.00; *P* =1).

**Figure 8.**
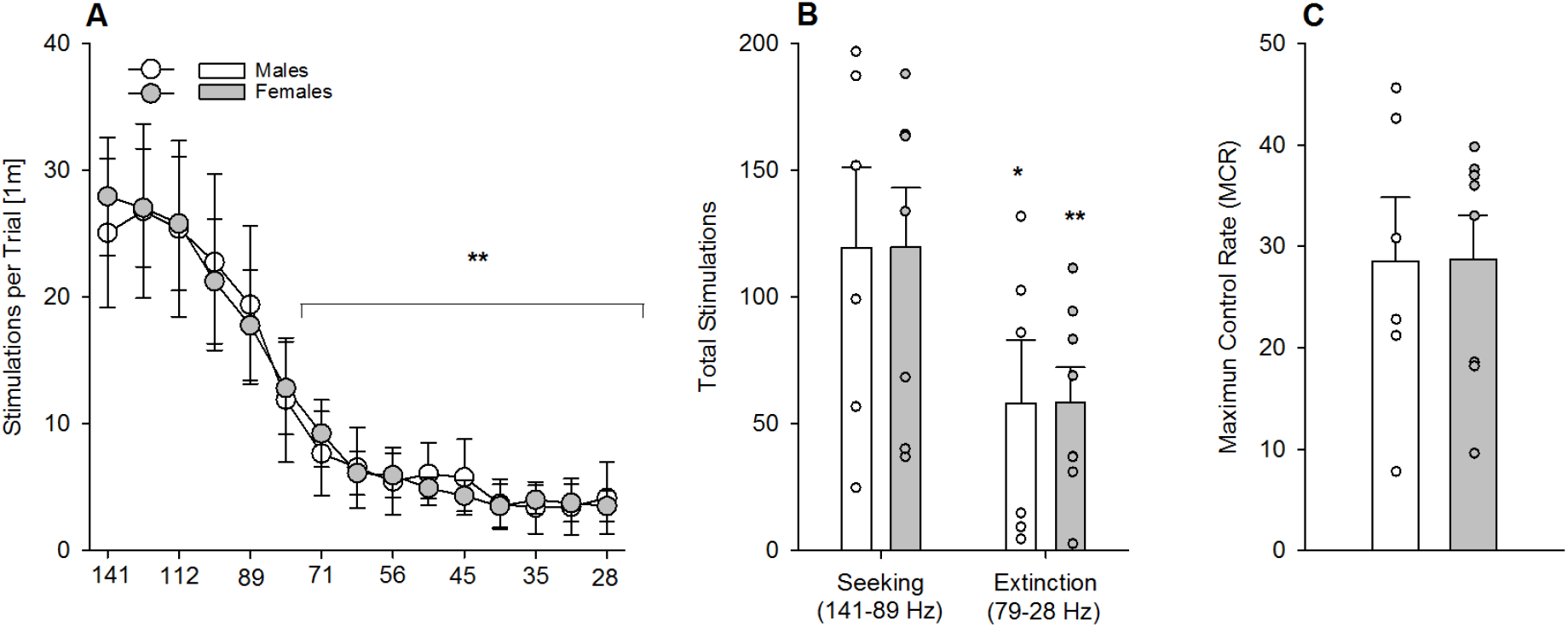
Characterization of brain reward sensitivity in male (n=6) and female (n=8) mice (Experiment 4). (A) Following ICSS training animals underwent the testing of 15 descending frequencies and all mice showed the expected frequency-dependent decrease in the number of stimulations per trial, which became significant from the 6^th^ frequency on. (B) Both seeking (first 5 frequencies) and extinction (remaining 10 frequencies) components were almost identical in all mice, which were characterized by a decrease in the extinction phase, compared to the seeking phase. (C) Likewise, all mice also showed an indistinguishable reinforcement rate. All data represent mean ± SEM. (*) and (**) indicate *P*<0.05 and *P*< 0.01 versus first 5 frequencies (A) or seeking (B), respectively.

## Discussion

We performed three experiments to examine and quantify addictive features of repeated excessive sugar consumption and one experiment to measure the reward sensitivity in male and female sugar-experienced C57BL6/N mice. In the first experiment, we show by means of thee DID procedure that female mice exhibited augmented binge-like intake of sugar as compared to male mice. In the second experiment, we show that following long-term free choice sugar consumption, female mice had a higher intake than male mice, especially during the active night phase. A deprivation phase resulted in an SDE that was again more pronounced in female mice and accompanied by a strong psychomotor stimulation. In the third experiment, we demonstrated enhanced persistency and motivation for sugar in female mice, but no sex difference in a compulsivity test. In experiment 4, we show that long-tern sugar consumption does not lead to sex-specific alteration in brain reward stimulation.

Using the DID procedure (Rhodes et al., 2005; Thiele and Navarro, 2014), sugar binging over a restricted 4 h period was induced in both sexes with a consumption of 50-75% more than the total amount consumed over continuous 24 h access. Although there is no consensus on the criteria defining a binge episode in rodents (i.e., hyperphagia, loss of control or/and duration of the event), it has been suggested that a two-fold increase of calorie intake relative to controls appears to be a reasonable criterion to determine hyperphagia (Valdivia et al., 2014); thus, mice in the DID procedure would fulfill the criteria of hyperphagia and show therefore sugar binge consumption. Females showed a more significant increase than males relative to continuous drinking, as well as higher binge drinking. Locomotor activity under continuous sugar exposure was higher in females, and interestingly, this difference was increased during binge conditions. This augmented activity can be seen as enhanced psychomotor stimulation, which is characteristic of a reward-seeking response (Sanchis-Segura and Spanagel, 2006).

We also studied long-term free choice sugar consumption. To date, chronic sugar consumption has almost exclusively been used to induce obesity in mice. In the diet/(sugar)-induced obesity model, the consequences on metabolic diseases or particular organ damage (i.e., liver) are of particular interest. However, the consequences of long-term free choice sugar consumption are usually not discussed in the context of addiction. Here we have followed our standard protocol of long-term ethanol drinking in mice (Eisenhardt et al., 2015b) and transferred it to voluntary sugar intake. As previously observed with alcohol, one key aspect involved in sugar drinking is the control by the circadian clock, resulting in diurnal rhythmicity of drinking behavior. Thus, regardless of short or long-term exposure, mice voluntarily consumed more sugar during the active, dark phase, than in the inactive, light phase. This diurnal rhythmicity remained stable over the entire time course of the experiment, including a deprivation phase. In comparison, following long-term alcohol or drug exposure and deprivation, disturbances in diurnal rhythmicity have been observed (Foo et al., 2017; Perreau-Lenz and Spanagel, 2015), which may be a distinction between sugar and drug addictive-like behavior.

After 6 weeks of long-term free choice sugar consumption, female mice showed approx. 50% more sugar consumption than males. These findings are consistent with what has been reported with alcohol in mice (Bilbao et al., 2019) and other drugs of abuse, in which female mice consume higher amounts of drugs under long access procedures (Lynch, 2018).

Following long-term free choice sugar consumption, a deprivation phase was introduced to induce relapse-like drinking referred to as the sugar deprivation effect (SDE) (Avena et al., 2005). Sugar deprivation for 2 weeks induced a robust SDE in all mice that was most pronounced during the first 4 hours after re-exposure to the sugar solution, similar to what we have previously found during the alcohol deprivation effect (ADE) in mice (Eisenhardt et al., 2015b). However, the SDE had a longer duration, as the increased intake persisted for at least two days, in contrast to the short-lasting ADE in mice (approx. 8 h). This suggests that in mice sugar deprivation may induce a higher “craving state” than alcohol. Furthermore, our data also indicate that females had a higher craving for sugar than males, as shown by the enhanced SDE. Moreover, during the second day, during which the intake slightly decreased in the males, the females had an almost identical consumption compared to the first SDE day. This phenotype seems to be specific for sugar, as we have previously shown that females do not show an ADE after a period of deprivation, probably due to high basal intake. Thus excessive baseline drinking usually results in decreased consumption, a phenomenon referred to as an inverse ADE (Vengeliene et al., 2014; Bilbao et al., 2019) that has been reported in various alcohol-preferring rat lines and high alcohol-preferring C57BL/6J mice (i.e., Vengeliene et al., 2003; Camp et al., 2011).

Another phenotype in female mice usually not observed with alcohol was a pronounced increase in locomotor activity during the SDE, which persisted up to 3 days. Thus, excessive sugar consumption can lead to enhanced psychomotor stimulation, a phenomenon seen after the intake of psychostimulants and in patients with attention deficit hyperactivity disorder (ADHD) caused by a disruption in dopamine signaling whereby dopamine D2 receptors are reduced in reward-related brain regions (Johnson et al., 2011). A similar mechanism may be in place in sugar binging, as rats show reduced mRNA levels for the D2 dopamine receptor in the brain reward system (Spangler et al., 2004).

We further applied operant sugar self-administration and quantified three criteria for addictive-like behavior – persistence, motivation and compulsivity. We found that female mice exhibited enhanced persistence and motivation – as measured by responding during the time out period and PR test, respectively – when compared to males. In contrast, there was no sex difference in compulsivity, as measured by quinine taste adulteration. These findings are consistent with rodent studies demonstrating that cue-reward learning in self-administration paradigms with drugs of abuse or natural rewards is more pronounced in females than in males (Lynch et al., 1999; Pitchers et al., 2015; Hammerslag and Gulley, 2014).

Finally, we used brain stimulation to determine whether the phenotype observed in females may be related to sugar-induced alterations in brain reward system sensitivity. Following long-term excessive sugar consumption, mice were tested for ICSS, which resulted in no difference between males and females. Thus, innate sex differences in the brain reward system do not contribute to the enhanced vulnerability for sugar in female mice, and long-term excessive sugar consumption in mice does not appear to alter reward processing. A potential limitation of the ICSS experiment was the lack of baseline testing of the mice prior to sugar exposure, which is a technically demanding experiment due to long-term electrode maintenance (8 weeks). Furthermore, since previous ICSS studies do not report any sex differences under naïve baseline conditions (Faunce and Banks, 2020; Lazenka et al., 2017), we found it reasonable not to test ICSS prior to sugar exposure.

In conclusion, our work suggests the occurrence of a sugar-related addictive-like phenotype in mice, similar but to some extent also different than that resulting from drugs of abuse (e.g., no alterations in diurnal rhythmicity following excessive sugar consumption and a longer duration of the sugar deprivation effect). Further studies are needed to determine whether these findings extend to different food components (i.e., sugary, salty and fatty foods) with respect to addictive-like behavior. Furthermore, we show enhanced vulnerability towards excessive sugar consumption, craving and relapse in female mice. Although more preclinical studies are needed, our finding of augmented excessive sugar consumption in female mice is consistent with a recent cross-sectional population-based study in more than 210.000 participants showing that women have higher intakes of sugar than men (Bennet et al., 2018). Whether women develop problematic sugar use more easily remains a matter of debate, but our study reinforces the idea that the occurrence of higher obesity rates in females may be due to the consumption of high sugar containing foods (Lovejoy and Sainsbury, 2009; Kanter and Caballero, 2012).

**Supported** by the BMBF (FKZ: 01ZX1909 - SysMedSUDs), and by the Deutsche Forschungsgemeinschaft (DFG) (TRR265 – Losing and Regaining Control over Drug Intake (Heinz et al., 2020).

## References

Avena NM, Hoebel BG. A diet promoting sugar dependency causes behavioral cross-sensitization to a low dose of amphetamine. Neuroscience 2003;122:17–20.

Avena NM, Long KA, Hoebel BG. Sugar-dependent rats show enhanced responding for sugar after abstinence: evidence of a sugar deprivation effect. Physiol Behav 2005;84:359–62.

Avena NM, Rada P, Hoebel BG. Sugar and fat bingeing have notable differences in addictive-like behavior. J Nutr 2009;139:623–8.

Becker JB, Koob GF. Sex Differences in Animal Models: Focus on Addiction. Pharmacol Rev 2016;68:242–63.

Becker JB, McClellan M, Reed BG. Sociocultural context for sex differences in addiction. Addict Biol 2016;21:1052–9.

Bennett E, Peters SAE, Woodward M. Sex differences in macronutrient intake and adherence to dietary recommendations: findings from the UK Biobank. BMJ Open 2018;8:e020017.

Bilbao A, Leixner S, Wei S, Cantacorps L, Valverde O, Spanagel R. Reduced sensitivity to ethanol and excessive drinking in a mouse model of neuropathic pain. Addict Biol 2019;24: 1008–1018.

Bilbao A, Neuhofer D, Sepers M, Wei SP, Eisenhardt M, Hertle S, Lassalle O, Ramos-Uriarte A, Puente N, Lerner R, Thomazeau A, Grandes P, Lutz B, Manzoni OJ, Spanagel R. Endocannabinoid LTD in Accumbal D1 Neurons Mediates Reward-Seeking Behavior. iScience 2020;23:100951

Bilbao A, Robinson JE, Heilig M, Malanga CJ, Spanagel R, Sommer WH, Thorsell A. A pharmacogenetic determinant of mu-opioid receptor antagonist effects on alcohol reward and consumption: evidence from humanized mice. Biol Psychiatry 2015;77:850–8.

Camp MC, Feyder M, Ihne J, Palachick B, Hurd B, Karlsson RM, Coronha B, Chen YC, Coba MP, Grant SG et al. A novel role for PSD-95 in mediating ethanol intoxication, drinking and place preference. Addict Biol 2011;16:428–39.

Cannella N, Cosa-Linan A, Büchler E, Falfan-Melgoza C, Weber-Fahr W, Spanagel R. In vivo structural imaging in rats reveals neuroanatomical correlates of behavioral subdimensions of cocaine addiction. Addict Biol 2018;23:182–195.

Cannella N, Halbout B, Uhrig S, Evrard L, Corsi M, Corti C, Deroche-Gamonet V, Hansson AC, Spanagel R. The mGluR2/3 agonist LY379268 induced anti-reinstatement effects in rats exhibiting addiction-like behavior. Neuropsychopharmacology 2013;38:2048–56.

Colantuoni C, Rada P, McCarthy J, Patten C, Avena NM, Chadeayne A, Hoebel BG. Evidence that intermittent, excessive sugar intake causes endogenous opioid dependence. Obes Res 2002;10:478–88.

Corwin RL, Grigson PS. Symposium overview-Food addiction: fact or fiction? J Nutr 2009;139:617–9.

Deroche-Gamonet V, Piazza PV. Psychobiology of cocaine addiction: Contribution of a multi-symptomatic animal model of loss of control. Neuropharmacology 2014;76 Pt B:437–49.

Eisenhardt M, Leixner S, Luján R, Spanagel R, Bilbao A. Glutamate Receptors within the Mesolimbic Dopamine System Mediate Alcohol Relapse Behavior. J Neurosci 2015a; 35:15523–38.

Eisenhardt M, Leixner S, Spanagel R, Bilbao A. Quantification of alcohol drinking patterns in mice. Addict Biol 2015b;20:1001–11.

Faunce KE, Banks ML. Effects of repeated kappa-opioid receptor agonist U-50488 treatment and subsequent termination on intracranial self-stimulation in male and female rats. Exp Clin Psychopharmacol 2020;28(1):44–54.

Foo JC, Noori HR, Yamaguchi I, Vengeliene V, Cosa-Linan A, Nakamura T, Morita K, Spanagel R, Yamamoto Y. Dynamical state transitions into addictive behaviour and their early-warning signals. Proc Biol Sci 2017;284:20170882.

Hammerslag LR, Gulley JM. Age and sex differences in reward behavior in adolescent and adult rats. Dev Psychobiol 2014;56:611–21.

Heinz A, Kiefer F, Smolka M, Endrass T, Beste C, Beck A, Liu S, Genauck A, Romund L, Banaschewski T, et al. Losing and regaining control over drug intake –from trajectories to mechanisms and interventions. Addict Biol 2020; 25(2):e12866.

Hudson A, Stamp JA. Ovarian hormones and propensity to drug relapse: a review. Neurosci Biobehav Rev 2011;35:427–36.

Johnson RJ, Gold MS, Johnson DR, Ishimoto T, Lanaspa MA, Zahniser NR, Avena NM. Attention-deficit/hyperactivity disorder: is it time to reappraise the role of sugar consumption? Postgrad Med 2011;123:39–49.

Kanter R, Caballero B. Global gender disparities in obesity: a review. Adv Nutr 2012;3:491–8.

Kendig MD. Cognitive and behavioural effects of sugar consumption in rodents. A review. Appetite 2014;80:41–54.

Lazenka MF, Suyama JA, Bauer CT, Banks ML, Negus SS. Sex differences in abuse-related neurochemical and behavioral effects of 3,4-methylenedioxymethamphetamine (MDMA) in rats. Pharmacol Biochem Behav 2017;152:52–60.

Lovejoy JC, Sainsbury A. Stock Conference 2008 Working Group. Sex differences in obesity and the regulation of energy homeostasis. Obes Rev 2009;10:154–67.

Lynch WJ, Carroll ME. Sex differences in the acquisition of intravenously self administered cocaine and heroin in rats. Psychopharmacology (Berl) 1999;144:77–82.

Lynch WJ. Modeling the development of drug addiction in male and female animals. Pharmacol Biochem Behav 2018;164:50–61.

Mancino S, Burokas A, Gutiérrez-Cuesta J, Gutiérrez-Martos M, Martín-García E, Pucci M, Falconi A, D’Addario C, Maccarrone M, Maldonado R. Epigenetic and Proteomic Expression Changes Promoted by Eating Addictive-Like Behavior. Neuropsychopharmacology 2015;40:2788–800.

Ogden CL, Carroll MD, Fryar CD, Flegal KM. Prevalence of obesity among adults and Youth: United States, 2011-2014. NCHS Data Brief. 2015 Nov;(219):1–8

Pelchat ML. Food addiction in humans. J Nutr. 2009;139(3):620–2.

Perreau-Lenz S, Spanagel R. Clock genes × stress × reward interactions in alcohol and substance use disorders. Alcohol 2015;49:351–7.

Pitchers KK, Flagel SB, O’Donnell EG, Woods LC, Sarter M, Robinson TE. Individual variation in the propensity to attribute incentive salience to a food cue: influence of sex. Behav. Brain Res 2015;278:462–9.

Reed GM, First MB, Kogan CS, Hyman SE, Gureje O, Gaebel W, Maj M, Stein DJ, Maercker A, Tyrer P, et al. Innovations and changes in the ICD-11 classification of mental, behavioural and neurodevelopmental disorders. World Psychiatry 2019;18:3–19.

Rhodes JS, Best K, Belknap JK, Finn DA, Crabbe JC. Evaluation of a simple model of ethanol drinking to intoxication in C57BL/6J mice. Physiol Behav 2005;84:53–63.

Sample CH, Davidson TL. Considering sex differences in the cognitive controls of feeding. Physiol Behav 2018;187:97–107.

Sanchis-Segura C, Becker JB. Why we should consider sex (and study sex differences) in addiction research. Addict Biol 2016;21:995–1006.

Sanchis-Segura C, Spanagel R. Behavioural assessment of drug reinforcement and addictive features in rodents: an overview. Addict Biol 2006;11:2–38.

Spanagel R. Animal models of addiction. Dialogues Clin Neurosci 2017;19:247–58.

Spangler R, Wittkowski KM, Goddard NL, Avena NM, Hoebel BG, Leibowitz SF. Opiate-like effects of sugar on gene expression in reward areas of the rat brain. Brain Res Mol Brain Res 2004;124:134–42.

Thiele TE, Navarro M. “Drinking in the dark” (DID) procedures: a model of binge-like ethanol drinking in non-dependent mice. Alcohol 2014;48:235–41.

Valdivia S, Patrone A, Reynaldo M, Perello M. Acute high fat diet consumption activates the mesolimbic circuit and requires orexin signaling in a mouse model. PLoSONE 2014;9:e87478.

Vengeliene V, Bilbao A, Spanagel R. The alcohol deprivation effect model for studying relapse behavior: a comparison between rats and mice. Alcohol 2014;48:313–20.

Vengeliene V, Siegmund S, Singer MV, Sinclair JD, Li TK, Spanagel R. A comparative study on alcohol-preferring rat lines: effects of deprivation and stress phases on voluntary alcohol intake. Alcohol Clin Exp Res 2003;27:1048–54.

Wiss DA, Avena N, Rada P. Sugar Addiction: From Evolution to Revolution. Front Psychiatry 2018;9:545.

